# Identification of LINE retrotransposons and long non-coding RNAs expressed in the octopus brain

**DOI:** 10.1101/2021.01.24.427974

**Authors:** Giuseppe Petrosino, Giovanna Ponte, Massimiliano Volpe, Ilaria Zarrella, Concetta Langella, Giulia Di Cristina, Sara Finaurini, Monia T. Russo, Swaraj Basu, Francesco Musacchia, Filomena Ristoratore, Dinko Pavlinic, Vladimir Benes, Maria I. Ferrante, Caroline Albertin, Oleg Simakov, Stefano Gustincich, Graziano Fiorito, Remo Sanges

**Affiliations:** Department of Biology and Evolution of Marine Organisms, Stazione Zoologica Anton Dohrn, Villa Comunale, 80121 Napoli (SZN), Italy; Neurobiology Sector, Scuola Internazionale Superiore di Studi Avanzati (SISSA), Via Bonomea 265, 34136 Trieste, Italy; Department of Integrative Marine Ecology, Stazione Zoologica Anton Dohrn, Villa Comunale, 80121 Napoli (SZN), Italy; Scientific Core Facilities & Technologies, GeneCore, European Molecular Biology Laboratory (EMBL), Meyerhofstrasse 1, 69117 Heidelberg, Germany; Marine Biological Laboratory (MBL), Woods Hole, Massachusetts, USA; Okinawa Institute of Science and Technology Graduate University, Onna, Okinawa 9040495, Japan; Department of Molecular Evolution and Development, Wien University, Althanstraße 14 (UZA I), 1090 Wien, Austria; Central RNA Laboratory, Istituto Italiano di Tecnologia (IIT), Via Enrico Melen 83, 16152 Genova, Italy

**Keywords:** Mollusks, Nervous system, Transcriptome, Transposable elements

## Abstract

**Background:** Transposable elements (TEs) widely contributed to the evolution of genomes allowing genomic innovations, generating germinal and somatic heterogeneity and giving birth to long non-coding RNAs (lncRNAs). These features have been associated to the evolution, functioning and complexity of the nervous system at such a level that somatic retrotransposition of long interspersed element (LINE) L1 has been proposed to be associated to human cognition. Among invertebrates, octopuses are fascinating animals whose nervous system reaches a high level of complexity achieving sophisticated cognitive abilities. The sequencing of the genome of the *Octopus bimaculoides* revealed a striking expansion of TEs which were proposed to have contributed to the evolution of its complex nervous system. We recently found a similar expansion also in the genome of *Octopus vulgaris*. However a specific search for the existence of full-length transpositionally competent TEs has not been performed in this genus.

**Results:** Here we report the identification of LINE elements competent for retrotransposition in *Octopus vulgaris* and *Octopus bimaculoides* and show evidence suggesting that they might be active driving germline polymorphisms among individuals and somatic polymorphisms in the brain. Transcription and translation measured for one of these elements resulted in specific signals in neurons belonging to areas associated with behavioral plasticity. We also report the transcription of thousands of lncRNAs and the pervasive inclusion of TE fragments in the transcriptomes of both *Octopus* species, further testifying the crucial activity of TEs in the evolution of the octopus genomes.

**Conclusions:** The neural transcriptome of the octopus shows the transcription of thousands of putative lncRNAs and of a full lenght LINE element belonging to the RTE class. We speculate that a convergent evolutionary process involving retrotransposons activity in the brain has been important for the evolution of sophisticated cognitive abilities in this genus.

## Background

Transposable elements (TEs) have contributed to the evolution of specific functions in a variety of biological systems and have given birth to a large fraction of vertebrate long non-coding RNAs (lncRNAs) [1–3]. Among TEs, retrotransposons move via a copy-and-paste mechanism using an RNA intermediate. The long interspersed element (LINE) L1, a non-LTR retrotransposon that predated the human genome, is active during neuronal differentiation [4] and causes somatic mosaicism establishing genomic variability in the brain [5]. Somatic L1 insertions are suggested to alter the transcriptional output of individual neurons, eventually affecting neuronal plasticity and behavior [6]. Transposition-driven genomic heterogeneity has also been documented in invertebrates including the neurons of mushrooms bodies of *Drosophila melanogaster* [7] where they have been suggested to drive behavioral variability in individual flies. Negative regulators of retrotransposons are reported to be expressed in specific subgroups of neurons in *Aplysia californica* [8], *Drosophila melanogaster* [7] and in the mouse brain [9], further supporting the idea that, in nervous systems, retrotransposons activity is finely regulated in a broad range of organisms and does not simply constitute noise. Our understanding of the activities of TEs in metazoa genomes is nevertheless still far to be complete and the number of cellular events in which they have an influence is constantly growing. Novel findings allow us to increase our comprehension but also add layers of complexity to the topic. L1 elements, for example, have been demonstrated to have specific activity also at the non-coding level in the regulation of transcription and the organization of the genome [10, 11] and to be a component of extra-chromosomal DNA [12].

Among invertebrates, the cephalopod mollusk *Octopus vulgaris* is known for the richness of its behavioral repertoire achieving sophisticated vertebrate-like plasticity and neural control [13–16]. The remarkable innovations in the morphological and functional organization of its nervous system [14, 17, 18] are linked to the evolution of an unprecedented complexity coded both at cellular and molecular levels [17–21]. In the common octopus, about 500 million nerve cells constitute the nervous system, with about 300 million composing the nervous system in the arm and about 200 million nerve cells in what is consider to be the central brain [22]. This number results to be ten thousand times higher than that found in the sea hare Aplysia, and still remains two hundred times higher when compared to the number of neurons present in the brain of the honeybee *Apis mellifera* (reviewed in Borrelli and Fiorito 2008). Octopus and their allies are also key examples of innovations in the evolution of invertebrates. Such innovations have been recognized not only at the level of the *Bauplan*, but include also extraordinary examples of the transcriptional and structural outputs of its genome such as: large cadherin genes encoding over 70 extracellular cadherin domains; unprecedented expansions of gene families crucial for regulation, signaling and cell comunication (e.g., protocadherins, zinc finger proteins, G-protein coupled receptors); birth of many novel octopus-specific genes; the existence of a vascular endothelial growth factor (VEGF) pathway; reflectin genes originated by horizontal gene transfer; differential arrangements of key developmental genes and extensive RNA editing capabilities [21, 24–28].

The sequencing of the *Octopus bimaculoides* genome [24] revealed an expansion of TEs and specific gene families related to transcriptional regulation and neuronal connectivity. Although no specific analysis was performed to identify full-length potentially active TEs and non-coding transcripts were discarded from the definition of the reference transcriptome, it was suggested that TEs are active in *O. bimaculoides* because a substantial fraction of RNAseq reads resulted in overlap with TEs fragments annotated in non-coding intergenic regions. Expansion of TEs has also been found in the genome of *Octopus minor* [29] and the other sequenced cephalopod species such as *Euprymna scolopes* [30] and the giant squid *Architeuthis dux* [31]. Performing a survey of the *O. vulgaris* genome we have confirmed the expansion of TEs also in this species [32]. The current picture is that, in cephalopod species, repeated elements cover on average 45% of the genome.

Here we sequence the *O. vulgaris* neural transcriptome to gain insights into the molecular composition underpinning its neural complexity. We identify a full-length LINE element and show that it is actively expressed in specific areas of the brain related to known forms of behavioral plasticity including learning. We also provide evidence for the transcription of thousands of long non-coding RNAs and the pervasive inclusion of TE fragments in coding and non-coding transcripts in both *O. vulgaris* and *O. bimaculoides*.

## Results

### Thousands of putative lncRNAs are expressed in the Octopus vulgaris nervous system

We generated a *de-novo* assembly of the *Octopus vulgaris* neural transcriptome identifying and evaluating its functional annotations, potential lncRNAs and repeats composition (Fig. 1; Supplemental Tables S1, S2; Supplemental Fig. S1, S2; Supplemental Files S1, S2, S3). From each of three different octopus individuals we collected four parts, three of them as representative of the central brain: 1) supra-esophageal mass (SEM), 2) sub-esophageal mass (SUB) and 3) optic lobe (OL); and one representing the peripheral nervous system: 4) a piece of the second left arm (ARM) including the arm nerve cord. We used these parts as source of RNA for the sequencing (see Methods, Supplemental Table S1). The sequencing generated approximately 850 million paired-end reads accounting for 85 Gbp of sequence data that, following assembly and filtering, produced 64,477 unique transcripts (Supplemental File S1). The sequences showed a N50 value of 2,087 bp, average length of 1,308 and about 38% average GC-content (Supplemental Table S2). The transcriptome resulted to be more than 98% complete and we functionally annotated 21,030 (32.6%) protein coding transcripts (Supplemental Fig. S1 and Supplemental File S3). By performing stringent annotation analysis a high proportion of transcripts (7,806; 12.1%) resulted to be putative lncRNAs (see Methods and Supplemental Fig. S2). In analogy to what is known about lncRNAs expression in mammals [33], the non-coding portion of the *O. vulgaris* transcriptome shows a lower level of expression when compared to protein coding genes (Fig. 1A) and a significantly higher number of lncRNAs results to be expressed in central brain (~10%) with respect to the arm (~7%, p-value < 1e-40) (Fig. 1B). Functional enrichments highlight differences between transcripts expressed in the central brain and in the periphery. Specifically, transcripts expressed in the brain result enriched in functions associated with neuronal cell adhesion and reverse transcription, while the ones expressed in the arm are enriched for functions associated with signal transduction and translation (Fig. 1C and Supplemental Fig. S3).

**Figure 1.**
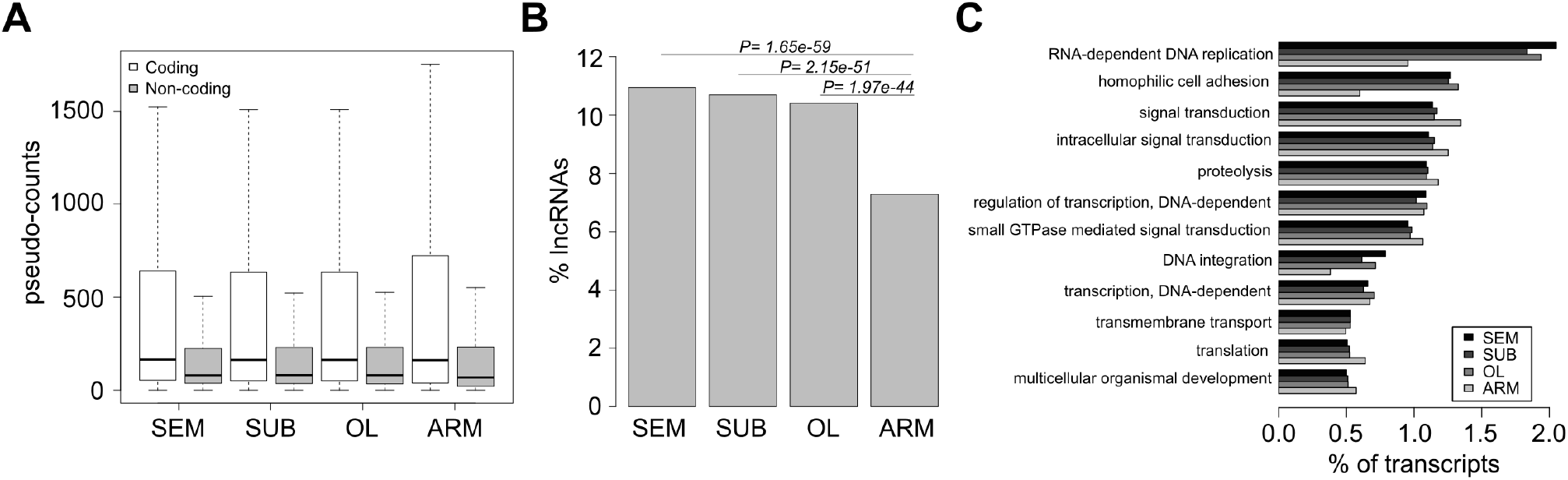
Features of the *Octopus vulgaris* brain and arm transcriptome. We sequenced the supra-esophageal (SEM) and sub-esophageal (SUB) masses and optic lobe (OL) as representatives of the brain, and the medial segment of an arm (ARM), including the arm nerve cord, as representative of peripheral system. (A) Expression levels for coding and non-coding transcripts. Non-coding transcripts are on average expressed at lower levels than coding. (B) Percentage of expressed non-coding transcripts. Brain samples result enriched for non-coding. (C) Percentages of transcripts expressed and their relative distribution among the most represented GO biological processes. A higher percentage of transcripts belonging to classes related to transposable elements and cell adhesion are expressed in the brain. Transcripts likely to be involved in signal transduction and translation constitute a larger quota in the arm.

We validated the expression and the sequence of selected coding and non-coding transcripts by RT-PCR and Sanger sequencing (Fig. 2) selecting a group of transcripts showing a specific peak of expression in each of the collected parts. A transcript was considered to have a peak of expression in a given part when showing an expression level higher than 0.5 counts per million (CPM) in all three biological replicates of exclusively one part and below 0.5 in all the others. This resulted in the selection of ~1,800 transcripts (~1,500 coding and ~300 non-coding). Among the coding transcripts with an expression peak only 54 resulted annotated. Among them we noticed the presence of putative homologs of homeobox genes and selected 4 of them for validation through RT-PCR in 3 different individuals. The tested *Arx* putative homolog (Aristaless Related Homeobox, comp31544_c0_seq1) resulted expressed mainly in the SEM by RNAseq and the RT-PCR validated this result. The RT-PCR also confirmed the peak of expression for the *Hoxb5a* putative homolog (Homeobox B5, comp28131_c1_seq2) which resulted expressed mainly in the SUB and the *Meox2* putative homolog (Mesenchyme Homeobox 2, comp34840_c15_seq1) which resulted expressed mainly in the ARM. The *Phox2b* putative homolog (Paired-Like Homeobox 2b, comp28142_c1_seq1) peaking in the OL by RNAseq data resulted expressed in all sampled parts of the brain by RT-PCR. For RT-PCRs concerning the lncRNAs, *Subl* (lncRNA with a peak of expression in the SUB, comp35227_c11_seq1) and *Arml* (lncRNA with a peak of expression in ARM, comp20195_c0_seq1) resulted to be tissue-specific, as they were mainly expressed in the SUB and in the ARM, respectively. On the other hand, *Seml* (lncRNAs with a peak of expression in SEM, comp18661_c0_seq1) resulted to be expressed in the SEM but also in the SUB and the OL, while *Oll* (lncRNA with a peak of expression in the OL, comp35506_c7_seq1) resulted expressed in the OL but also in all the other sampled parts (Figure 2C). Both *Oll* and *Arml* show the existence of two different isoforms in at least one individual. The RT-PCR results were generally in agreement with the sequencing data for both coding and non-coding transcripts tested. The identifiers indicated for every transcript are the same used in the Supplementary files and can be used to identify the corresponding sequences and annotations.

**Figure 2.**
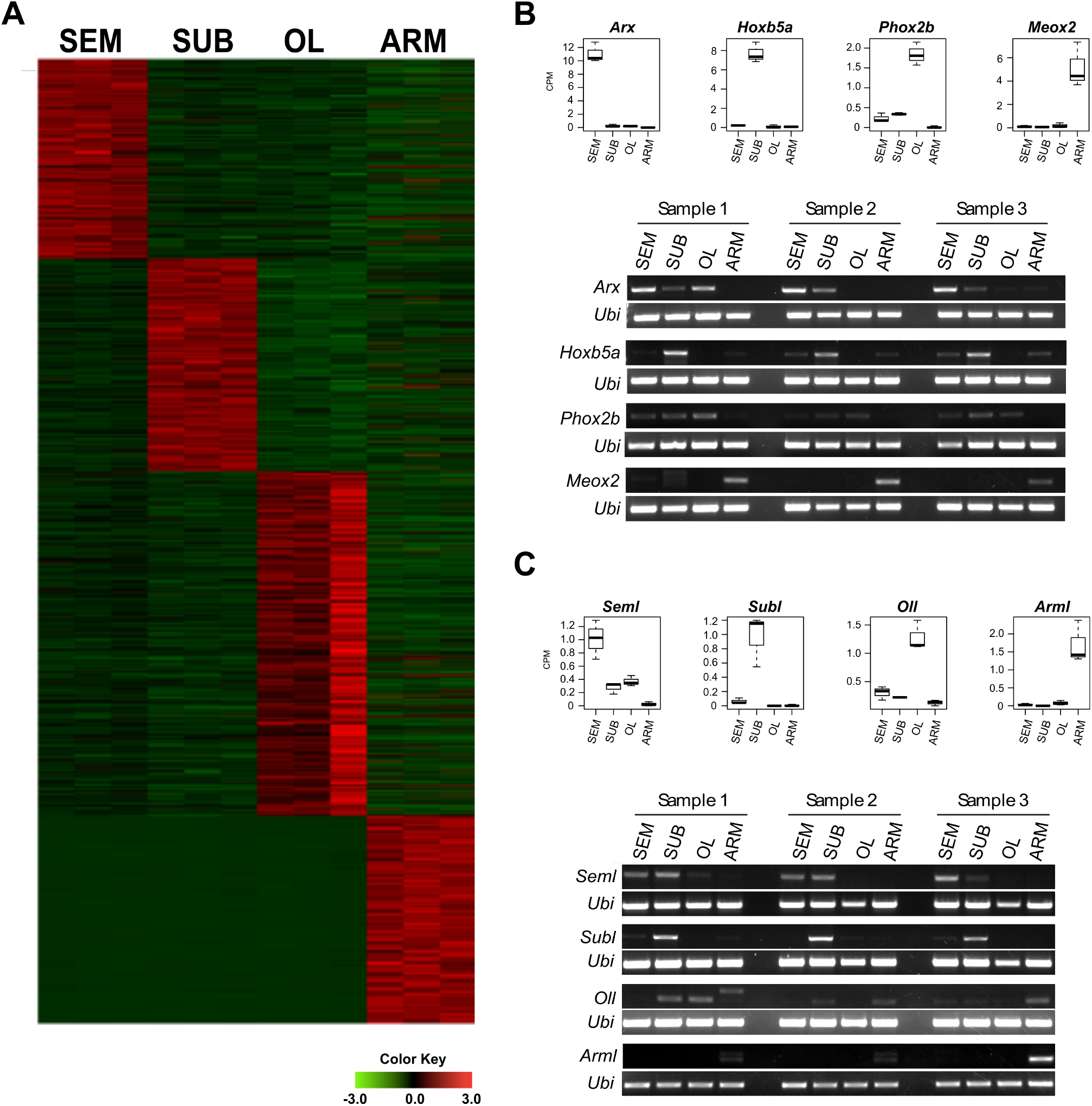
Transcripts with an expression peak and transcriptome validations. (A) Heatmap showing the expression levels for transcripts classified as having a peak of expression. (B) Boxplots showing RNAseq expression levels of coding transcripts selected for validation and their relative RT-PCR results from 3 different individuals. (C) Boxplots showing RNAseq expression levels of non-coding transcripts selected for validation and their relative RT-PCR results from 3 different individuals. The octopus ubiquitin transcript (Ubi) has been used as positive control in all the experiments.

In order to verify the presence of lncRNAs also in *O. bimaculoides* we reassembled the public RNAseq data from this species with the same method used for our *O. vulgaris* data (see Methods). We assembled 92,820 unique transcripts (Supplemental Table S2 and Supplemental File S4) of which 84,043 (90%) map on the published assembled genome with at least 90% coverage and 90% identity (Supplemental File S5). Our analysis demonstrated the presence, also in this octopus species, of thousands of putative lncRNAs (File S6). Indeed we were able to classify 10,030 assembled transcript as putative non-coding. They correspond to more than 10% of the assembled transcriptome of which 9,132 map on the reference genome.

### A full-length LINE element is transcribed in the Octopus vulgaris brain

In order to evaluate the contribution of TEs to the *O. vulgaris* transcriptome we analyzed its content in terms of repeated elements using RepeatMasker (Supplemental File S7) and found that more than 3.5 millions of nucleotides derive from interspersed repeats (4.5% of the total transcriptome content; Supplemental Table S3). More than 35% of the generated transcripts contain at least one interspersed repeat fragment. Among them, retroelements represent the TE fragments more frequently embedded in transcripts (26% of transcripts contain a fragment from at least one retroelement). According to the segregation of transposable elements in transcripts expressed in the different sequenced parts of the organism, we observed that SINEs fragments are present in a higher fraction of transcripts expressed in the brain, while LINEs, LTRs and DNA transposons are present in a higher portion of transcripts expressed in the periphery (Supplemental Fig. S4A). SINE also results the class of retroelements more frequently embedded in lncRNAs (Supplemental Fig. S4B).

To identify the molecular basis of the observed TEs expansion in *O. vulgaris* we searched the transcriptome for putative autonomous active elements. We found a single element, a LINE mainly transcribed in the brain (Fig. 3A-C) that we named RTE-2_OV following phylogenetic analysis from which the element resulted to belong to the RTE clade (Fig. 3D). The identified LINE presents an ORF of 3,327 nucleotides and 5’ and 3’ UTRs of about 600 nucleotides. The translation corresponds to a 1,109 aminoacids polypeptide chain containing all the catalytic aminoacids and domains needed for retrotransposition (Fig. 3A,B): a C-terminal endonuclease (EN), a reverse transcriptase (RT) and an N-terminal C2H2 zinc-finger (Znf) which is relatively rare in RTE elements [34]. The finding that this element contains an intact ORF and is transcribed indicates that it might be active and possibly drive retrotransposition. We then asked whether this element might show evidence of somatic retrotransposition. Southern blots analysis did not lead to conclusive results because of high background noise likely due to the high number of copies of this TE in the genome of *O. vulgaris*, however the pattern observed is consistent with the existence of germinal and somatic genomic variations associated to the element (Supplemental Fig. S5). We also performed quantitative PCR experiments on DNA deriving from the four parts of different individuals. Results add support to the presence of somatic retrotransposition in the brain (Fig. 3E) showing an average higher content of RTE-2_OV DNA in the SUB with respect to the other parts. This result suggests the existence of a higher amount of RTE-2_OV DNA in cells from the SUB, whereas cells from other parts show a similar average content. The existence of these increased amounts of RTE-2_OV DNA in SUB is in line with its higher expression in the SUB (Fig. 3F). Interestingly, the putative homolog of Piwi, known to repress the translation of TEs and therefore to restrict retrotransposition, displays a lower expression in the SUB with respect to the other portions of the central brain of (Fig. 3G) consistently with what has been observed in the fruitfly brain [7].

**Figure 3.**
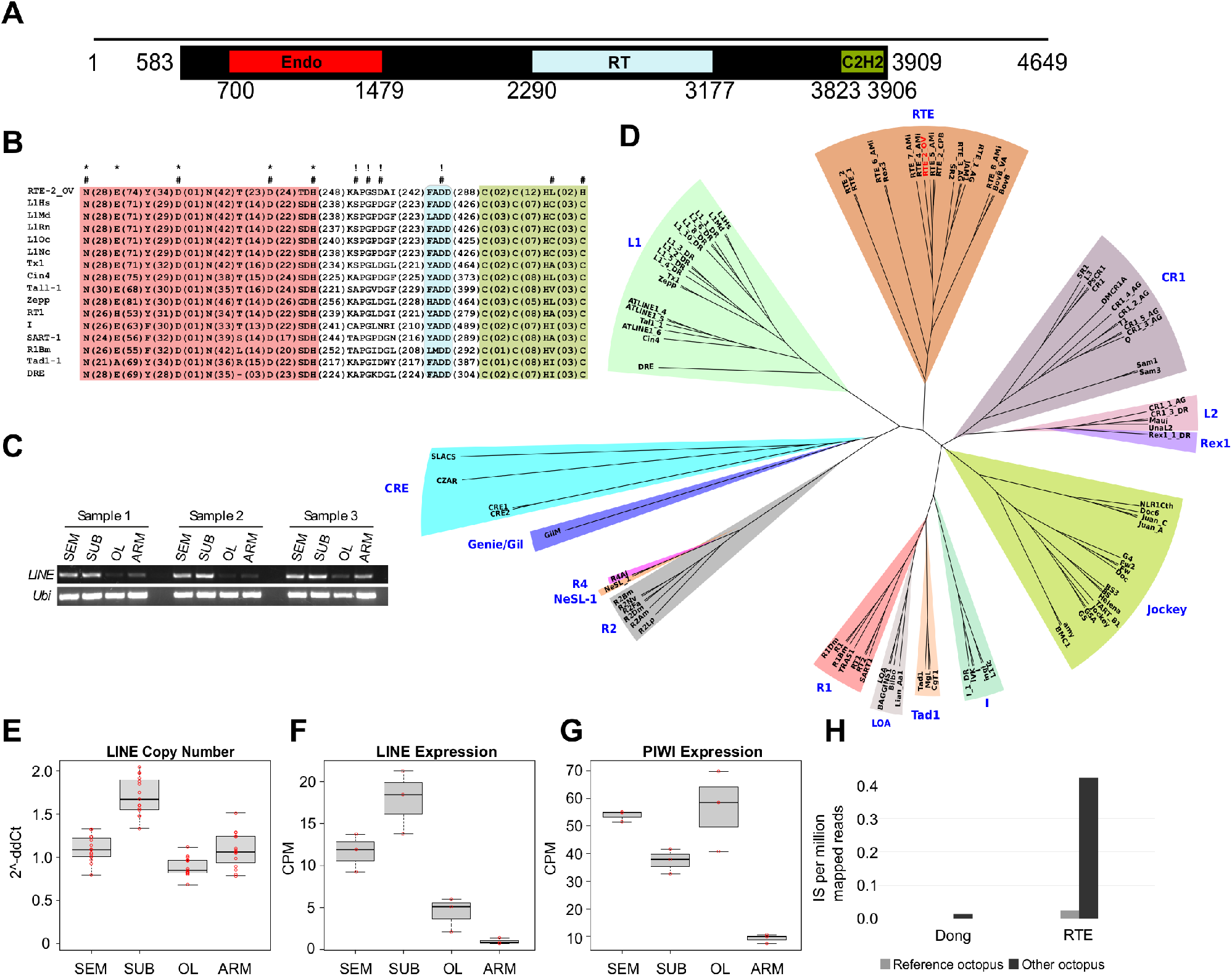
A full-length potentially active LINE transcribed in *Octopus vulgaris*. (A) Domain composition of the discovered LINE (comp36575_c1_seq3). Black line represents the transcript, black box the location of the ORF and colored boxes the protein domains (Endo, endonuclease; RT, reverse transcriptase; C2H2, zinc finger). Numbers are relative to nucleotide positions in the transcript. (B) Schematic alignment highlighting the conserved catalytic aminoacids in the group of LINEs adapted from [37] plus the octopus element. Color code of the domains is the same as in A; aminoacids critical for EN (*), RT (!) and retrotrasposition (#). (C) Electrophoresis from RT-PCR of the LINE showing expression from three different animals. (D) Phylogenetic tree based on 100 LINEs from [38] (see Supplementary Table 4) plus the octopus element in red. (E) LINE copy number variation analysis using quantitative real-time PCR with Taqman probes. (F) Expression levels of the LINE based on RNAseq data. (G) Expression levels of Piwi-like protein 1 (comp33731_c0_seq1) from RNAseq data. (H) Normalized number of non-reference insertions in two different *O. bimaculoides* individuals (octopus used for the reference genome and a different octopus individual) for the two LINEs identified in this species.

In order to add further support to the existence of active LINEs in the genomes of the Octopus *genus* we searched the public genome, transcriptome and our custom transcriptome assembly (Supplementary Files S4-S6) of *O. bimaculoides* for the presence of assembled full-length LINEs. We were not able to identify any assembled full-length LINE in the transcriptome nor in the genome. However, when inspecting the repeat library consensus generated in the work by Albertin *et al*. [24] we identified two LINEs with a full-length ORF and the complete set of domains: an RTE and a Dong [35] element. To gain insights into the possibility that the identified LINE elements might be active we generated DNAseq sequencing data from the optical lobe of an *O. bimaculoides* individual and searched for novel integration sites (IS) with respect to the reference genome using MELT [36]. The same analysis was performed using the public DNAseq data derived by the gonads of the individual used to assemble the reference *O. bimaculoides* genome. Comparing the whole genome sequencing data from the two different *O. bimaculoides* individuals we obtained significant evidences of activity only for the RTE element. Indeed, the element showed a significantly higher number of non-reference ISs with respect to the reference individual (Fig. 3H, p-value 4.8e-69) which indicates that this element is at the basis of polymorphisms between the two individuals. Conversely, the Dong element resulted in a very low number of non-reference ISs which does not significantly differ from the results obtained with reads from the reference individual and therefore does not display evidences of recent activity.

### The RTE-2_OVelement is mainly expressed in amacrine neurons

To identify the domains of activity for the RTE-2_OV element we performed RNA *in-situ* hybridization and immunohistochemistry analysis. Localization of the RTE-2_OV transcript in *O. vulgaris* through *in-situ* hybridization (ISH) showed specific expression of the element in subgroups of neurons in brain and absence of any signal in neuropils. We found most of the small cells of the sub-frontal lobe and of the five gyri of the vertical lobe as the most intensely stained areas in the supra-esophageal mass (SEM; Fig 4A-E; Supplemental Fig. S6B-E). Neural cells stained for RTE-2_OV create a tapestry closely matching the pattern obtained by DAPI (Supplemental Fig. S6B,C; Supplemental Fig. S6D,E as control), indicating that the transcript is expressed in the great majority of the cellular bodies. We also found several positive neural cells in the sub-esophageal mass (SUB: anterior and posterior areas; Fig. 4F-H; Supplemental Fig. S6G) and in the optic-lobe (OL; Fig. 4I-K; Supplemental Fig. S6L). In particular after ISH, positive cells (20-25 μm in diameter) were observed at the level of the SUB in the pallovisceral lobe, and some larger neurons (40-50 μm) belonging to typical cellular-types of the motor-centre present in the area. Positivity to the mRNA of the RTE-2_OV was also observed in cells belonging to the ventral side of the anterior pedal lobe (SUB) and in some larger cells (up to 50 μm) at the level of the dorsal brachial lobe (anterior part of the SUB). The small cells pertaining to these areas do not show any positive signal. In the optic lobe, the outer layer appeared rich of intensely positive cells (small amacrine cells, < 5 μm) and the inner medulla presented scattered cell bodies (up to 10 μm) expressing the element. Cell bodies of the peduncle complex at the level of the median and posterior lobules of the olfactory lobe also revealed a positive signal after ISH. We finally found isolated sparse large motor neurons positive at the level of the nerve cord in the arm (Supplemental Fig. S6S). No positive cells were observed in any muscle fiber in the arm or any other structure.

**Figure 4.**
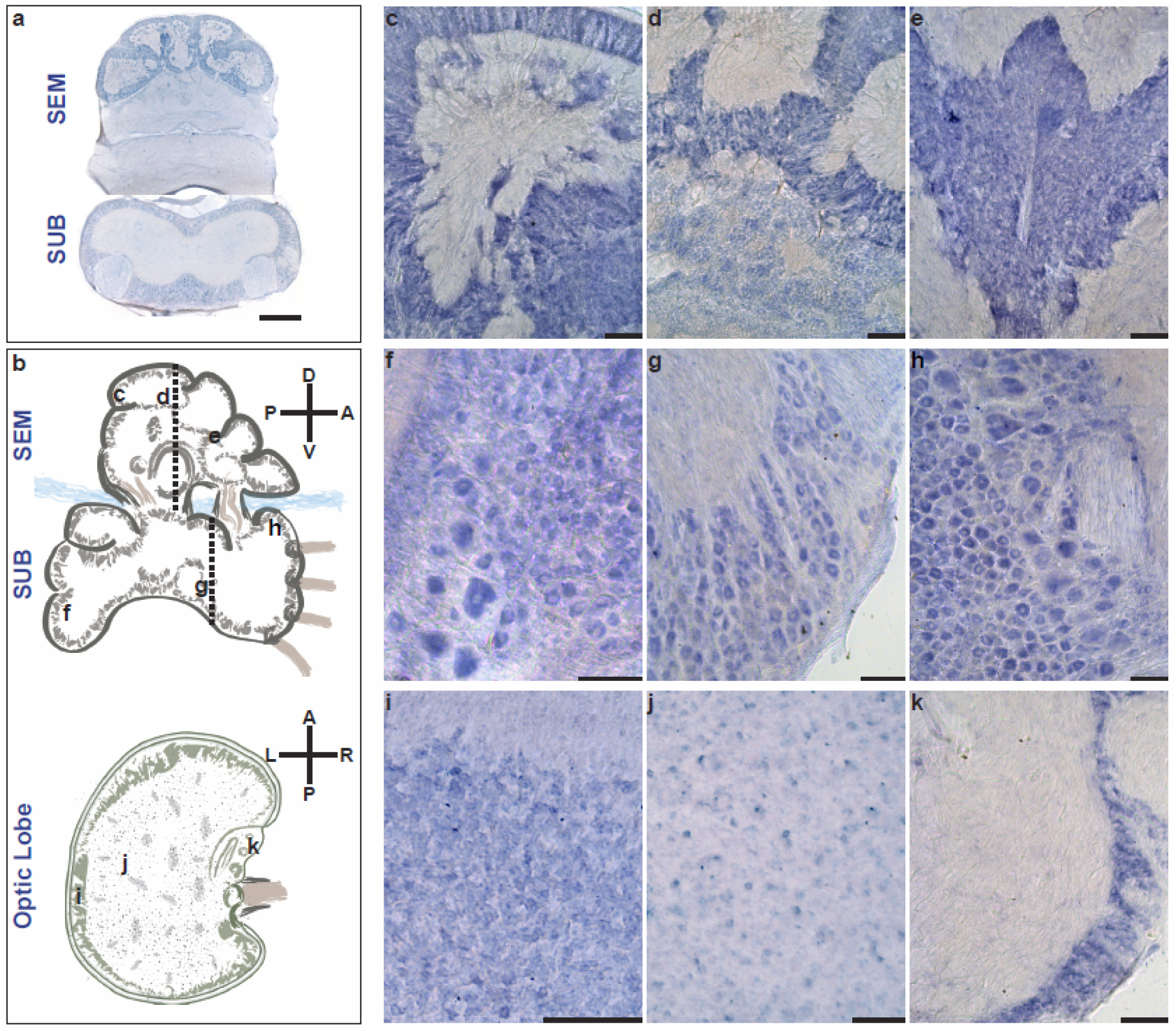
Localization of RTE-2_OV mRNA by in situ hybridization. (A) Bright-field micrographs of coronal sections of the supraesophageal (SEM) and subesophageal (SUB) masses hybridized to digoxigenin-labelled LINE antisense (refer to the plane indicated by the dashed-lines in B). The five gyri of the vertical lobe (dorsal in SEM) appear positively marked by the RTE-2_OV mRNA. Several cells in the cortical layers of the SUB appear also stained after in situ hybridization. Neuropil of SEM and SUB do not show signal. (B) Schematic outline of the parts of the octopus brain for SEM and SUB (sagittal plane), and the optic lobe (OL; horizontal plane). Axes illustrating dorso-ventral and antero-posterior (SEM and SUB), and antero-posterior and left-right (OL) orientations with respect to the octopus body plan. Black letters indicate approximate levels of the sections provided in the other panels of the figure. (C) Detail of a gyrus of the vertical lobe (SEM) with densely packed amacrine cells showing positive signal. (D) A similar signal in the gyri of the vertical lobe and some scattered positive cells in the sub-vertical lobe. (E) Section at the level of the sub-frontal lobe with densely packed amacrine small cells showing strong positive signal. In the SUB we observed in (F) positive cells (20-25 μm in diameter) in the pallovisceral lobe and some larger neurons (40-50 μm) belonging to typical motor-centre cellular types. (G) Cells (20-25 μm) belonging to the ventral side of the anterior pedal lobe, and at the level of the dorsal brachial lobe (H) where some larger cells (up to 50 μm) are also marked after ISH. The small cells pertaining to these areas do not show positivity. Details of horizontal sections of the *O. vulgaris* optic lobe (I, J, K; in areas indicated in B): (I) Outer layer rich of intensely positive cells (small amacrine cells, <5 μm); (J) Inner medulla with scattered LINE mRNA-expressing cell bodies (up to 10 μm); (K) cell bodies of the peduncle complex at the level of the median and posterior lobules of the olfactory lobe (cells of about 10 μm). Scale bars: 100 μm; 500 μm in a. Schematic drawings in (B) not to scale.

Immunohistochemistry (IHC) with RTE-2_OV custom antibodies identified in the vertical lobe a number of large cells organized in trunks and positive fibers in the neuropil of the gyri. We also observed large positive cells organized in chain at the level of the cellular wall of the sub-vertical lobe (Fig 5B) and few positive neurons in the great majority of the islands of cells in the posterior wall of the sub-vertical lobe. A distinct pattern of positive fibers was found in neuropils of the SEM (Fig. 5A-C) together with a scattered number of cell bodies (see areas belonging to superior- and inferior frontal lobes; Fig. 5C-D). We did not identify positive fibers in the SUB, but only a distinct population of a small number of neurons in the vasomotor lobes (Fig 5E) and in discrete areas of the anterior and the lateral posterior wall of the pedal lobe (Fig. 5F-H). Immuno-reactivity was also evident in several scattered amacrine cells of the external granular layer of the OL (Fig 5I). We found several RTE-2_OV positive fibers dispersed in discrete areas of the neuropil of the OL and a distinct pattern of positive cells and fibers in the peduncle lobe, mostly towards the internal layer of cells of the neural wall of the olfactory lobe and the spine (Fig. 5L). A small number of large cells dispersed in the arm nerve cord were also found to be stained (data not shown).

**Figure 5.**
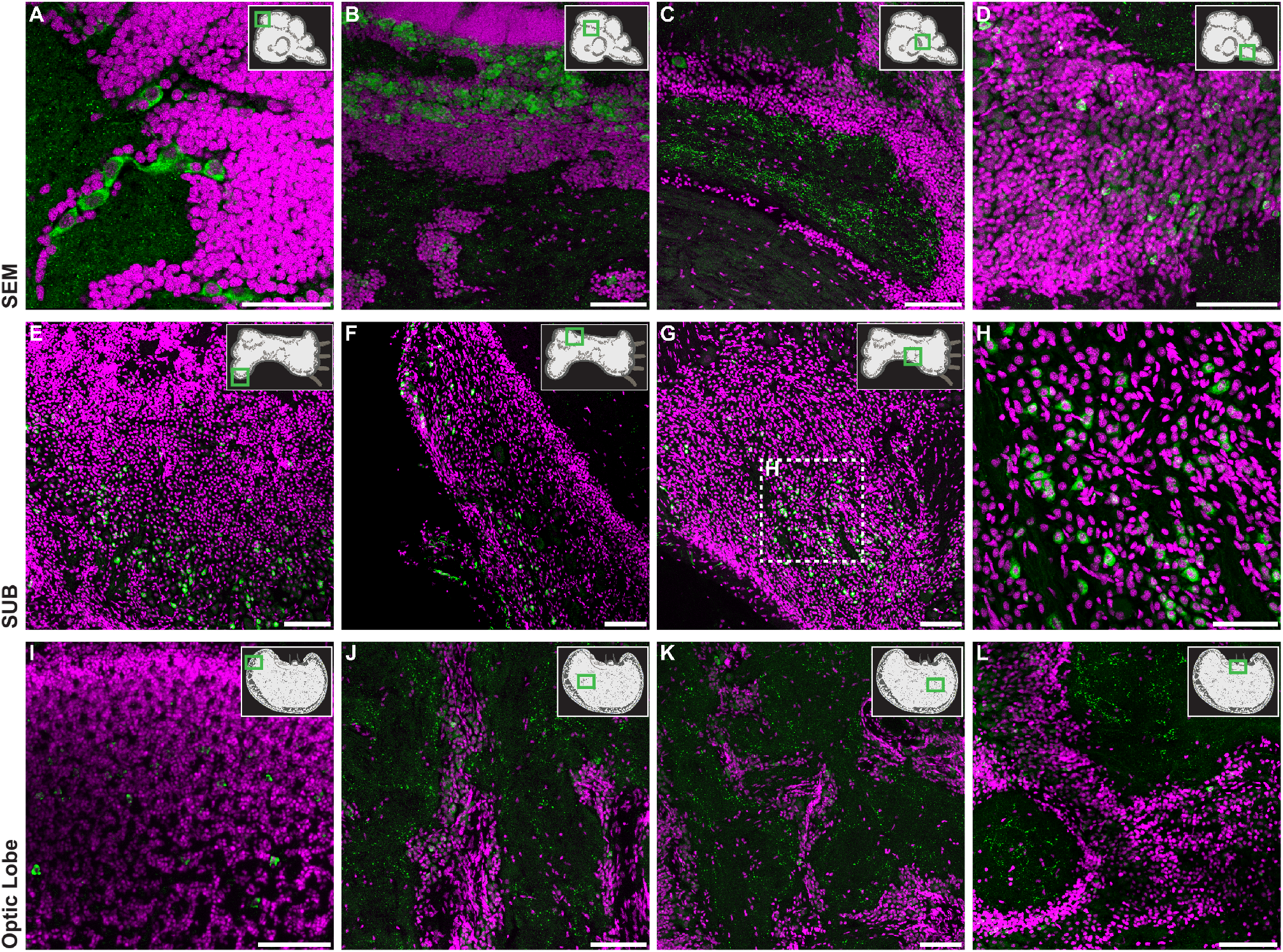
RTE-2_OV immunostaining in different areas of the brain. Coronal sections of the supra- (SEM; A-D) and sub-oesophageal (SUB; E-H) masses, and horizontal sections for the optic lobe (OL; I-L) following fluorescent-IHC (RTE-2_OV signal in green, DAPI used as nuclear stain in magenta) highlight a differential pattern of positive cells and fibers in *O. vulgaris* brain. A schematic drawing of the brain parts is provided with areas of interest indicated in the green square. (A) Large positive cells are found in the vertical lobe (VL). These appear organized in trunks and clearly distinguishable from the population of numerous amacrine cells constituting the VL (DAPI stained layer). (B) Large cells in the sub-vertical lobe (cellular wall), and a part of the bundle of fibers present at the beginning of the sub-frontal lobe (C). (D) Scattered positive cells are also identified in the posterior buccal lobe. Several positive cells are identified in the SUB in the cellular walls of the vasomotor lobe (E) and in discrete areas of the pedal lobe (F). A similar pattern of positive cells is recognized at the level of the anterior part of the pedal lobe (G). A detail in the higher magnification (H; square in G) of the cellular layer of the lobe serves to highlight the population of positive cells. In the OL several amacrine cells are found positive in the external granular layer (I). The OL-medulla is populated by few immune-reactive neurons found in the cellular islands (J, K), and positive fibers dispersed in the surrounding neuropil (J, K). Positive cells and fibres are also identified in the peduncle lobe. The internal layer of cells of the neural wall of the olfactory lobe and the spine (L) are shown. Scale bars: 100μm, with the exception of A and H (50μm).

## Discussion

The analysis of the coding part of the *O. bimaculoides* genome provided interesting parallelisms with the transcriptional output of mammalian genomes [24]. Our results in *O. vulgaris* support and expand this view suggesting that the molecular organization of the octopus neural transcriptome resembles the mammalian one also for what concerns the transcription of thousands of lncRNAs and their TEs content.

Here we report the assembly of the most complete transcriptome for the *Octopus vulgaris*. The sequencing of different portion of the organism allowed us to identify transcripts whose expression showed a peak in only one part. For PCR validations we chose putative homologs of homeobox genes. *Arx* is an homeobox gene associated with mental retardation. In mammalian it has been shown to be expressed in fetal brain at various developmental stages, mainly in neuronal precursors in the telencephalon were it results involved in corticogenesis and in the differentiation and maintenance of specific neuronal cell types [39, 40]. In our *O. vulgaris* transcriptome assembly its putative homolog results to be expressed mainly in the SEM consistently with an important function in the central brain also in this species. As another example, the expression of the putative homolog of *Meox2* appears to be restricted to the arm in *O. vulgaris*, which also appears consistent with the known function of this gene in vertebrates were it controls limb muscle development [41]. The selection and validation of these expression peaking transcripts together with the functional information of the putative homologs added support to the quality of the assembled transcriptome and allowed us to perform more specific analyses. We observed that more than 10% of the assembled transcriptome for *O. vulgaris* and *O. bimaculoides* is composed of putative lncRNAs and that more than 35% of the assembled transcripts contain at least one fragment derived from a transposable element. Despite the fact that different classes of TEs appear to be differentially segregated between coding and non-coding transcripts we did not find an overall difference nor a strong enrichment for lncRNAs to contain embedded TE fragments with respect to coding ones. This result was unexpected and warrant further investigation. One of the causes might rely in our conservative strategy to call non-coding transcripts for which we might likely have underestimated the fraction of lncRNAs increasing the rate of false negative.

The observation reported from us and others about TEs expansion in octopus drove our interest toward the proportion of the transcriptome deriving from transposable elements. This lead us to the discovery of a novel RTE LINE retrotransposon transcribed in the central nervous system of *O. vulgaris*. Reanalysis of the *O. bimaculoides* data allowed us to identify, also in this species, a LINE member of the RTE class showing evidences of activity. We could not find any full length copy of the *O. vulgaris* element in the genome survey [32] nor could we find the *O. bimaculoides* element full length sequence in its reference assembly. This likely resulted from the difficulties to assemble a reference genome using exclusively short reads in regions containing long repeated sequences as those containing full-length LINE elements. Despite this technical gap, the complementary use of transcriptomic, repeats reconstruction, annotation data and manual curation allowed us to identify the full length autonomous elements likely to be active in both the octopus species. The identification of potential integration polymorphisms of the RTE element in *O. bimaculoides* is a strong sign of its activity. On the other hand, the results from quantitative PCR and Southern blot analysis relative to the element discovered in *O. vulgaris*, although not conclusive, support the existence of somatic retrotransposition in the central brain for this species. It is here important to consider that results from quantitative PCR for LINEs genomic copy number variations should be interpreted with care, especially in case of absence of additional supporting evidences. Indeed, in addition to a potential variation in genomic copy number, they might also reflect a different amount of extra-chromosomal DNA which, at least in human, is composed by a consistent fraction of LINEs DNA [12, 42]. More specific and careful inspections are needed to validate, beyond any reasonable doubt, the activity of the elements presented in this work. Nevertheless our results represent the first evidences of LINE activity conserved in the Octopus genus.

Through ISH we localized RTE-2_OV transcripts in specific groups of neural cells and areas of the *O. vulgaris* nervous system, with no signal in neuropils. In particular, positive signals were observed at the level of the majority of amacrine cells (about 3 micron diameter) which are suggested to have very limited neural processes [43]. The most intriguing signals are those present at the level of the sub-frontal lobe and of the five gyri of the vertical lobe (VL). These areas appeared to be the most intensely positive to RTE-2_OV transcript found in the supra-esophageal mass. VL represents about 14% of the overall volume of the SEM in the adult *O. vulgaris* [44] and counts about 25 million neural cells, equivalent to more than 65% of the total number of nerve cells estimated for the SEM in the common octopus [22]. The largest quota of this impressive number of closely packed nerve cells is represented by the amacrine cells, the smallest in the octopus brain. Lying below the vertical lobe is the sub-vertical lobe. Its neural architecture is characterized by the presence of a wall that in several regions is folded to form islands of cells. Many cells have a diameter less than 5 μm, and very few are larger than 10 μm. At both sides of the sub-vertical lobe there are cells with diameter of 5-10 μm, but also the largest neurons of the whole SEM reaching 25 μm. Through IHC in these areas we identified a number of large cells positive to RTE-2_OV antibodies appearing organized in trunks as typically described by Young in 1971 [45]. We also observed large positive cells organized in chains at the level of the cellular wall of the sub-vertical lobe, and few positive neurons in the great majority of the islands of cells (posterior wall of the sub-vertical lobe). Finally, immuno-reactivity was also evident in several scattered amacrine cells belonging to the external granular layer of the OL suggested to correspond to the neural cells described to project in the underlying plexiform zone [15].

Remarkably, *O. vulgaris* posterior frontal and vertical lobes belong to a neural circuit considered to be functionally analogous to the mammalian hippocampus and limbic lobe [15, 17, 46, 47] where retrotransposons are also reported to be active [7, 48]. The VL is also suggested to be analogous to the mushroom bodies of insects [49]. All other octopus brain areas where the mRNA and/or the protein were identified are centers of active neural plasticity for visual sensory-motor processing and memory (SEM and OL), and motor processing (SUB) [17, 45, 50]. The positive cells overlap with the intricate neural network made by amacrine cells known to act as amplifying matrices functioning as read-in read-out memory of both sensory-motor visual and chemo-tactile processing [15, 47, 51, 52].

The reported imaging experiments allow also to made observations at the cellular level. The signals resulting from IHC and ISH are principally cytoplasmic. An higher number of cells appear to contain the element mRNAs with respect to the cells in which the protein can be identified. We only observed a partial overlap between cells expressing the mRNA and cells containing the protein. These observations might reflect the complexity of LINEs involvement in cellular activities. Such activities might be resulting by the strategies employed in the cell to contrast retrotransposition but they can also reflect the basis of specific LINE functions that we currently understand only with limitations. The most of our knowledge on LINE elements comes from studies on human L1. Its transcript is a bicistronic mRNA which codifies for two open reading frames (ORF1 and ORF2). Among them, the ORF2 protein contains the endonuclease, reverse transcriptase and zinc finger domains and therefore it can be considered the ORF codifying for the retrotranspositional machinery and the functional homolog of the single ORF of the RTE elements that we found in the two octopus species. Of note, the two human L1 ORFs do not result to be translated at the same levels. ORF1 is translated at a higher level than ORF2, and different proteins which repress ORF2 translation have recently been identified [53]. Consequently, the ORF2 protein has been observed to be absent in a portion of cells where the mRNA and ORF1 protein were present [54]. The limited efficiency for the translation of ORF2 has been proposed to be one of the mechanisms that the cell puts in action to limit and/or regulate the retrotranspositional potential of the L1 mRNAs. Indeed, for the retrotransposition to take place, all the L1 components, ORF1 protein, ORF2 protein and L1 mRNA must be present in the same ribonucleoprotein complex (RBP) within the nucleus. The lack of any of these components in the RBP does not allow retrotransposition. Reasoning along this line, another mechanism used to restrict retrotransposition is the confinement of the RBP complex within the cytoplasm, where it is assembled. As long as the complex is located in the cytoplasm it cannot access the genome and therefore cannot retrotranspose. For this reason, cells subjected to retrotransposition have been identified mainly in cells which underwent duplication. The mechanism proposed is that the RBP can get in contact with nuclear DNA only after the nuclear membrane dissolves during mitosis. If the RBP remains in contact with the DNA it is then found into the nucleus after the nuclear membrane of the new cell has been rebuilt thus allowing retrotransposition [54, 55].

Spatial uncoupling between L1 mRNA and proteins can also be observed independently from cellular defense mechanisms. Indeed, L1 elements are also active at the mRNA level as lncRNAs. During early development and in the regulation of cell potency, L1 transcription and its mRNA result to play a crucial role in regulating global chromatin accessibility and the L1 mRNA can act as a nuclear RNA scaffold to recruit specific regulatory proteins to the genome [10, 11]. Another molecular event resulting in spatial uncoupling of LINEs transcription/translation can be observed when LINE mRNA is used from a given cell type to target a different cell type with retrotranspositional activity. In fruitfly the existence of a mechanism producing mRNAs of the I element (another member of the LINE class) by the nurse cells has recently been observed. Once produced, these mRNAs are transported to the transcriptionally inactive oocytes were they are translated and can cause retrotransposition (Wang et al. 2018).

## Conclusions

Our data corroborate the hypothesis that LINE elements, potentially capable to determine genomic and functional diversity among individual cells, might be active and functionally important in the central nervous system of highly intelligent organisms such as octopus [6]. The localization of RTE-2_OV mRNA into different neural cells and not in the neuropil or muscle, complemented by the distinctive pattern of expression of its protein product and the identification of a potentially active element of the same class from a congeneric species, support the view that RTE-2_OV might be functional in neural centers in *O. vulgaris* and make it an important candidate to be further studied for its contribution to neural plasticity in this fascinating organism.

## Methods

### Animals, sampling and ethical statement

*Octopus vulgaris* were collected by local fishermen from the Bay of Naples (Southern Tyrrhenian Sea, Italy) in early summer 2012. Animals were transported to the Stazione Zoologica Anton Dohrn in Napoli and maintained according to a standardized acclimatization protocol [57–59]. Samples were taken from local fishermen, by applying humane killing following principles detailed in Annex IV of Directive 2010/63/EU as described in the Guidelines on the Care and Welfare of Cephalopods [59] and following protocol for collection of tissues described by Baldascino and coworkers [60]. Death was confirmed by transection of dorsal aorta. All dissections were carried out on seawater ice bed. During surgery optic lobes (OL), supra-(SEM) and sub-esophageal (SUB) masses were dissected out from the animal, a piece of an arm (ARM), usually the second left arm, was also taken. The complete dissection lasted less than 10 min. Sampling from live animals occurred before the entry into force of the Directive 2010/63/EU in Member States and therefore no legislation was in place in Italy regulating research involving cephalopods. However, the care and welfare of animals has been consistent with best practice [59, 61, 62] and in compliance with the requirements of the Directive 2010/63/EU that includes cephalopods within the list of species regulated for scientific research involving living animals. In addition, animals killed solely for tissue removal do not require authorization from the National Competent Authority under Directive 2010/63/EU and its transposition into national legislation.

### RNA extraction and sequencing

For each octopus (N=3) total RNA was isolated from central nervous tissues (SEM, SUB and OL) and ARM, a part of the body including the largest quota of the neuronal population belonging to the peripheral nervous system [22, 63] and thus constituted by muscle and peripheral nervous tissue. SV total RNA isolation kit (Promega, #Z3100) was utilized according to the manufacturer’s protocol. DNA was degraded by treating samples with Turbo DNase Kit (Ambion) according to the manual. Quality and quantity of RNA was assessed by NanoDrop (Thermo-Fisher) and RNA BioAnalyzer (Agilent Technologies, Santa Clara, CA, USA). Paired-end libraries were prepared using the Illumina TruSeq RNA sample library preparation kit (Illumina, San Diego, CA, USA). Each sample was barcoded, samples pooled and sequenced in two lanes on the Illumina HiSeq 2000 platform (paired-end, non-strand specific, 2×50bp read length protocol).

### Raw reads quality filtering

Quality of raw reads was assessed using FastQC (release 0.10.1). Raw reads were filtered and trimmed based on quality and adapter inclusion using Trimmomatic [64] (release 0.22; parameters: -threads 24, -phred 64, ILLUMINACLIP:illumina_adapters.fa:2:40:15, LEADING:3, TRAILING:3, SLIDINGWINDOW:3:20, MINLEN:25). Read pairs with both reads passing the filters were considered for the transcriptome assembly. Trimmed and filtered reads were normalized to remove duplicates using the normalize_by_kmer_coverage.pl script from Trinity [65] (release r2013_08_14; parameters: --seqType fq, --JM 240G, --max_cov 30, -- JELLY_CPU 24).

### De novo assembly and quantification of transcript abundances

Transcriptome was assembled using Trinity (release r2013_08_14) on the trimmed, filtered and normalized reads exploiting the Jaccard clip to limit assembly of chimeras. Assembly was performed using the following parameters: --seqType fq, --JM 240G, --inchworm_cpu 24, --bflyHeapSpaceInit 24G, --bflyHeapSpaceMax 240G, --bflyCalculateCPU, --CPU 24, --jaccard_clip, --min_kmer_cov 2. To measure expression levels, raw reads were mapped on the assembled transcriptome using Bowtie (version 1; parameters: -p 24, --chunkmbs 10240, --maxins 500, --trim5 2, --trim3 2, --seedlen 15, --tryhard -a -S). SAM outputs from Bowtie [66] were converted into BAM, sorted, indexed and counted using the view, sort, index and idxstats programs from Samtools [67]. All transcripts not showing at least 0.5 reads mapping per million mapped reads (CPM) in at least 2 samples were discarded from the transcriptome as being expressed at too low levels and therefore likely deriving by noise or assembly artifacts.

### Annotation of the assembled transcriptome

CEGMA (Core Eukaryotic Genes Mapping Approach; release 2.5) [68] was used to measure the completeness of the assembled transcriptomes using the set of 248 Core Eukaryotic Genes (CEGs). Transcripts annotation was performed using the Annocript pipeline [69] (release 0.2) with the combination of tool, parameters and databases described below and using BLAST+ (release 2.2.27) [70]. To annotate proteins we used BLASTX against the UniRef90 and Swiss-Prot databases from UniProt (release 2013_09) [71] with parameters: -word_size 4, -evalue 10e-5, -num_descriptions 5, -num_alignments 5, -threshold 18. To annotate protein domains we used RPSBLAST against the Conserved Domains Database (CDD v3.10) [72] with parameters: -evalue 10e-5, -num_descriptions 20, -num_alignments 20). Ribosomal and small noncoding RNAs were identified using BLASTN against a custom database made by Rfam (realease 11.0) [73] and ribosomal RNA sequences from GenBank (parameters: -evalue = 10e-5, -num_descriptions 1, - num_alignments 1). Each transcript was associated to Gene Ontology (GO) [74], Enzyme Commission (EC) [75] and UniPathway [76] through cross-mapping of the best match from UniRef90 or Swiss-Prot using the annotation mapping tables from UniProt. For each transcript, we used the Virtual Ribosome (Dna2pep release 1.1) [77] to predict the length of the longest ORF searching across all reading frames without the constraint to begin translation from a methionine start codon (parameters: -o none, -r all). The non-coding potential for each transcript was calculated using Portrait (release 1.1) [78].

### Non-coding annotation of the assembled transcriptome

Putative lncRNAs were classified based on a heuristic process considering the annotation results. The constraints used to identify potential lncRNAs have to be considered very stringent (Extended Data Fig. 2). In published studies, different combinations of analyses have been used to identify lncRNAs [78–81]: 1) lack of similarity with proteins, 2) lack of similarity with domain profiles, 3) lack of similarity with other RNAs (ribosomal, snoRNA, etc.), 4) transcript and longest ORF lengths, 5) non-coding potential. We put all these together and classified as lncRNA only those transcripts satisfying all the following conditions: a. length >= 200 nucleotides, b. lack of similarity with any of the following: protein from Swiss-Prot and UniRef90, domains from CDD, rRNA from GenBank and other small ncRNA from Rfam, c. longest ORF < 100 aminoacids, d. non-coding potential score >= 0.95. Using these stringent constraints, we were able to predict in the *O. vulgaris* transcriptome 7,806 (~12%) transcripts as putative lncRNAs.

### Assembly, mapping and annotation of the Octopus bimaculoides public RNAseq data

*O. bimaculoides* RNAseq raw data from Albertin et al. [24] were downloaded from NCBI SRA in October 2015 using the SRA Toolkit. Raw reads were filtered and trimmed based on quality and adapter inclusion using Trimmomatic (release 0.33; parameters: -threads 32, ILLUMINACLIP:illumina_adapters.fa:2:40:15:10:true LEADING:3 TRAILING:3 SLIDINGWINDOW:3:20 MINLEN:50). Read pairs with both reads passing the filters were considered for the assembly. Trimmed and filtered reads were assembled with Trinity (release 2.1.0; parameters: --seqType fq --SS_lib_type RF --CPU 32 --max_memory 240G --inchworm_cpu 32 --bflyHeapSpaceInit 24G -- bflyHeapSpaceMax 240G --bflyCalculateCPU --normalize_reads --min_kmer_cov 2 --jaccard_clip) using digital normalization, strand information, the Jaccard clip and assuring that every kmer used in the assembly was present in at least 2 reads to reduce noise. Redundancy of assembled transcripts was reduced using Cd-hit [82] (version: 4.6, parameters: -c 0.90 -n 8 -r 0 -M 0 -T 0). To measure the expression levels, raw reads were mapped on the transcriptome using Bowtie (version 1, parameters: -t -q -p 32 --chunkmbs 10240 -maxins 500 --trim5 2 --trim3 2 --seedlen 28 --tryhard -a -S). SAM outputs from Bowtie were converted into BAM, sorted, indexed and counted using the view, sort, index and idxstats programs from the Samtools software collection. All transcripts not showing at least 1 reads mapping per million mapped reads (CPM) in at least 1 sample were discarded from the transcriptome. *Octopus bimaculoides* genome was downloaded on August 2015 from http://genome.jgi.doe.gov/pages/dynamicOrganismDownload.jsf?organism=Metazome. Assembled and filtered unique transcripts were mapped on the genome using gmap [83] (version: 2015-09-28, parameters: --suboptimal-score 0 -f gff3_gene --gff3-add-separators 0 -t 32 --min-trimmed-coverage 0.9 --min-identity 0.9) considering only transcripts aligning at least 90% of their length with 90% minimum identity. We were able to map 84,043 (~90%) transcripts. Annotations and all the remaining analysis were executed as for *O. vulgaris*.

### Identification and classification of repetitive elements

Repetitive elements for each transcriptome were annotated using RepeatMasker (A.F.A. Smit, R. Hubley & P Green RepeatMasker at http://repeatmasker.org; release 4.0.5) searching against the Repbase database [84] (release 20140131) with parameters: -species bilateria, -s, -gff. We counted from RepeatMasker output the repeat fragments present at least once in each transcript and built a matrix containing the percentage of transcripts containing fragments related to a. retroelements, b. DNA transposons, c. satellites, d. simplerepeats, e. low complexity, f. other, and g. unknown classes for each transcriptome according to the RepeatMasker classification.

### Identification of full-length, putatively active transposable elements

We parsed RepeatMasker output calculating the percentage of overlap between the assembled transcripts and the repeat consensus from Repbase selecting all elements with at least 80% coverage on the repeat consensus. Elements showing the highest coverage were selected. On these we used Virtual Ribosome to predict the longest complete ORFs by searching across all reading frames with methionine as start codon and a canonical stop (parameters: -o strict, -r all). A single transcript resulted with a complete ORF. On this we used InterPro [85] to identify and classify protein domains. The potential catalytic amino-acids essential for the retrotransposition were manually identified comparing the putative translation with those reported in Clements and Singer [37]. The same analysis was performed on both the transcriptomes of *O. bimaculoides* (assembled by Albertin and assembled by us). The analysis was also performed on the RepeatMasker annotation of the genome by Albertin downloaded from http://octopus.unit.oist.jp/OCTDATA/. For consistency, we also analyzed RepeatMasker annotations of the genome and the transcriptomes produced using the same tool, library and parameters used for *O. vulgaris* and the other species considered in this study. In no-one of the analyses we could find a full-length transposable element retaining a complete ORF for *O. bimaculoides*. We then translated the main RepeatScout and RepeatModeler repeat libraries consensuses assembled by Albertin et al. (main RepeatScout library: http://octopus.unit.oist.jp/OCTDATA/TE_FILES/mainrepeatlib.fa.gz; RepeatModeler library: http://octopus.unit.oist.jp/OCTDATA/TE_FILES/oct.rm.tar.gz) with the Virtual Ribosome tool to predict longest ORF searching across all reading frames showing methionine as start codon (parameters: -o strict, -r all) and a canonical stop. The InterPro tool was then used to identify and classify the LINE characteristic domains. The potential catalytic amino-acids essential for the retrotransposon activity were manually identified comparing the ORF sequences with those reported in Clements and Singer. This led us to the identification of two potentially functional LINE retrotransposons.

### Identification of retrotrasposition events in O. bimaculoides

The two candidates retrotransposons found in *O. bimaculoides* RepeatModeler libraries were analyzed to search for integration sites in gonads and optic lobe using MELT [36] (v2.0.2). Two genomic DNAseq WGS libraries from gonads (SRR2010220 and SRR2005790) were downloaded from the European Nucleotide Archive (ENA) at https://www.ebi.ac.uk/ena/data/view/PRJNA270931. We generated two additional DNAseq WGS libraries from DNA extracted from the optic lobe (L001 and L002) of a different individual. *O. bimaculoides* reference genome was filtered for scaffold shorter than 10,000 bp and reads mapped on it using BWA [86] (v0.7.15; parameters: mem, -t 32). SAM output from BWA were converted into BAM, sorted, indexed and counted using the view, sort, index and idxstats programs from the SAMtools software. The resulted sorted BAM files were used as input for MELT (parameters: -d 10000). Since the reads length differ between the two set of libraries (150 bp for gonads and 260 bp for optic lobe) the optic lobe dataset was trimmed with Trimmomatic (v0.32; parameters: CROP:200, HEADCROP:50) to obtain homogeneous reads of 150 bp in all the datasets. We filtered the integration sites (ISs) identified by MELT for entries which passed all the MELT checks and which presented at least 3 discordant pairs of reads supporting both left and right sides of the breakpoints. BLAST (v2.6.0; parameters: -evalue 99999) search of the candidate retrotransposons consensus sequences was performed against the genome and the identified ISs were additionally filtered out when the BLAST search showed similarity in a range of 260 bp around the IS breakpoint. The same analysis was performed using non-trimmed reads and two additional ISs identification programs, RetroSeq [87] and an in-house developed pipeline, and the significance of results was maintained (data not shown).

### Phylogenetic tree generation

Evolutionary tree in Figure 3D was generated using 100 full-length LINEs belonging to 15 LINE clades (Extended Data Table 5). Protein sequences were selected from Ohshima and Okada [38] and manually checked. InterPro was used to identify endonuclease and reverse transcriptase domains in all the LINEs. Multiple sequence alignments were performed using MAFFT [88] (v7.221; with option L-INS-i). We utilized TrimAl [89] (v1.4.rev15) to perform automated trimming aligned sequences (parameters: -fasta - automated1). Phylogenetic relationships between LINE elements was reconstructed with MrBayes [90] (v3.2.1). Bayesian analysis was run for six million generations with twenty-two chains, sampling every 1,000 generations (6,000 samples). Convergence was attained with standard deviation of split frequencies below 0.01 and a consensus tree was generated using a burnin parameter of 1,500 (25% of 6,000 samples). The phylogenetic tree was visualized with FigTree program (release 1.4.2; http://tree.bio.ed.ac.uk/software/figtree/).

### Classification of transcripts expressed in each sample, expression peaks and selection of candidates for validations

We classified subsets of transcripts according to their expression levels across the different parts. A transcript was considered expressed in a specific part if, in all the replicates of that part, it showed an expression level > 0.5 CPM. This resulted in the classification of the groups of transcripts used to perform the analysis (see: Figure 1c, Extended Data Fig. 4). We also classified transcripts having a peak of expression. These represent transcripts showing an expression level > 0.5 CPM in all three biological replicates of exclusively one part and below 0.5 in the others, which resulted in ~1,800 transcripts. They were used to draw the heatmap (Extended Data Fig. 3a) using MeV (release 4.8.1) as part of the TM4 suite [91] with hierarchical clustering exploiting Pearson correlation. Candidates for validations were selected among the 1,800 transcripts with expression peak using the following additional criteria. For coding transcripts, we randomly selected four coding transcripts representing octopus homologs of homeobox genes. The annotation was manually verified. For lncRNAs we randomly selected four putative non-coding containing a SINE fragment. Both coding and lncRNA candidates were validated using RT-PCR and Sanger sequencing.

### Polymerase chain reaction (PCR)

Octopus cDNAs were generated from 200 ng of total RNA using Superscript VILO cDNA Synthesis Kit (Life Technologies) in 20 μl reaction volume. PCR were carried out using 20 ng of cDNA, 0.25 μl of Taq DNA Polymerase (5U/μl; Roche), 1 μl of each specific forward and reverse primer (25pmol/μl), 2.5 μl of PCR reaction buffer (10x), 2.5 μl of dNTP mix (10x) and water (up to final volume: 25 μl). The Ubiquitin gene (accession number FJ617440) was used as internal control. Reactions for the coding transcripts were amplified with a single step of 2 min at 94°C, 15 s at 94°C, 30s at 60°C and 1 min at 72°C for 35 cycles, and 7 min for 72°C. Reactions for the non-coding transcripts were amplified at annealing temperature of 58°C. The following primers were utilized:

Arx forward 5’-TCCCTGCCTTCTCAACACAT-3’
Arx reverse 5’-TCCGAACTTCCACGCTTACT-3’
Hoxb5a forward 5’-GTGGCGAGGAATTTAGGAAG-3’
Hoxb5a reverse 5’-GCAACAGTCATAGTCCGAACAG-3’
Phox2b forward 5’-AATGGGGTGAGATCCTTTCC-3’
Phox2b reverse 5’-TTCATTGCAATCTCCTCTCG-3’
Meox2 forward 5’-TCCAGAACCGTCGGATGAAA-3’
Meox2 reverse 5’-TACGTAAAGGGCACACACCT-3’
Seml forward 5’-CACTTGTGCAAGGTACCACG-3’
Seml reverse 5’-AGGTCTCCTTAAATTTATTTCTGTGCA-3’
Subl forward 5’-ACAGAGCATCTTGAGTCTCACT-3’
Subl reverse 5’-CACTCCTGCGCCTTTCATTT-3’
Oll forward 5’-GGATTGACCCTGCAACTTGG-3’
Oll reverse 5’-CAGTGATGACGGACTTGCAA-3’
Arml forward 5’-GTACCCCACAAAATTAAATC-3’
Arml reverse 5’-CACTCACAAGGCTTTAGTTGGC-3’
Ubi forward 5’-TGTCAAGGCAAAGATTCAAGA-3’
Ubi reverse 5’-GGCCATAAACACACCAGCTC-3’

### LINE copy number variation using quantitative real-time PCR

Copy number variation analysis was performed on genomic DNA extracted from octopuses (N=9; SEM, SUB, OL and ARM). One ARM sample was chosen as calibrator, while 18S was chosen as invariant control. Purified genomic DNA concentrations were assessed by NanoDROP (ThermoFisher Scientific). According to the starting concentration, DNA samples were diluted in TE buffer (10mM Tris-HCl, 1mM EDTA, pH 7.5) to a concentration of 100 ng/μL and then further diluted to a concentration of 10 ng/μL. All dilutions were checked by NanoDROP. Primers and multiplexing efficiencies were verified by linear regression to a standard curve ranging from 50 ng to 16 pg of genomic DNA. LINE and 18S slopes were −3.3 and 3.8 respectively, and represent acceptable amplification efficiencies. Standard curves also confirmed that the final concentration of 5ng DNA tested in qPCR was within the linear range of reaction. Reactions were performed in 20 μL reaction mixture containing iQ Multiplex Powermix (Bio-Rad), Taqman primers (10 μM) and probes (10 μM) differentially labeled (with FAM or VIC fluorophore) and specifically designed to hybridize with the target DNA sequences. LINE element was amplified using the following primers:

LINE forward 5’-AGCAGTGGGAATCATTCA-3’
LINE reverse 5’-GTCGTTTTCGTCGAACCAGT-3’
18S was amplified using the following primers:
18S forward 5’-AGTTCCGACCGTAAACGATG-3’
18S reverse 5’-CCCTTCCGTCAATTCCTTTA-3’
As probe sequences we utilized:
FAM: 5’-AACTCTGGGCCAAATTACGA-3’
VIC: 5’-GGGAAACCATAGTCGGTTCC-3’

qPCR was carried out for 20s at 90°C, followed by 40 cycles of 10s at 95°C and 30s at 59°C using 7900HT Fast Real Time PCR System (Applied Biosystems). Assays were performed for each sample in duplicate and reproduced four times. Data obtained from the co-amplifications of the target DNA sequence and the internal invariable control 18S were analyzed using the 2–ΔΔCt method79.

### Southern Blotting

The brains (SEM, SUB and OL) and a piece of an arm (ARM) of three *O. vulgaris* were dissected after humane-killing and immediately frozen in liquid nitrogen. Pulverized samples were treated following the methods utilized by Perelman and coworkers80; in brief, after phenol ∶chloroform(50:50)extractionDNA chloroform (50:50) extraction DNA was precipitated using cold isopropanol followed by centrifugation, suspended in TE buffer (10 mM Tris– HCl pH 8.0 and 0.1 mM EDTA pH 8.0), treated with ribonuclease A (10 μg /ml) and incubated at 37°C for 30 min. DNA concentration was estimated using NanoDrop and quality checked by electrophoresis on 0.8% agarose gel. 10 μg genomic DNA for each tissue was digested with EcoRI (New England Biolabs) overnight at 37°C and resolved on a 0.9 % agarose gel for 15 hours at 1.5V. DNA was transferred to a Hybond-N+ nylon membrane (0.45 μm; Amersham Pharmacia Biotech) according to Sambrook and Russell81. DIG-labeled LINE DNA probe was prepared by PCR DIG Probe synthesis kit (Roche). Hybridization and autoradiography were performed according to the DIG Application Manual (Roche).

### Probe synthesis for in-situ hybridization

We amplified by PCR a 356-bp cDNA fragment of the assembled LINE from bp 1,512 to bp 1,868 using the following primers:

LINE forward 5’-GGGTCAGAAAGTGACGAGGA-3’
LINE reverse 5’-TGCATGAGGCGGAGTTTAG-3’

The choice of the fragment and the design of primers have been based on manual curation steps ensuring that the chosen fragment is present exclusively in the transcript of the identified LINE element and in no other assembled transcripts. The amplified fragment was cloned into TOPO^®^ TA Cloning^®^ vector (Life Technologies, CA, USA) according to manufacturer’s protocol. Cloned fragment was digested using BAMHI and ECORV restriction enzymes and validated by Sanger sequencing. Sense and antisense digoxigenin-labelled RNA probes were generated by in vitro transcription using the DIG-RNA Labeling Kit (SP6/T7; Roche Applied Sciences, QC, Canada). Labeled RNA probes were quantified by dot blot analysis.

### In-situ hybridization experiments

Brain masses (SEM, SUB, OL) and a segment of an arm from octopuses (N = 3) were fixed in paraformaldehyde 4% (PFA) in phosphate buffered saline (PBS) at 4°C (3h for brain masses; overnight, ARM). Samples were washed (four rinses in PBS), dehydrated in series of graded methanol/PBS (1:3, 1:1, 3:1 v/v) and stored at least one night in methanol (−20°C). Tissues were then rehydrated at 25°C in a series of graded methanol/PBS (3:1, 1:1, 1:3 v/v) solutions and cryoprotected in 30% sucrose in PBS. After sucrose infiltration, samples were embedded in tissue freezing medium (OCT; Leica Biosystems) and sectioned using a cryostat (Leica CM3050 S). Sagittal and/or coronal sections (40 μm) were collected in PBST (phosphate buffered saline including 0.1% Tween™ 20 and 0.2mM sodium azide). Washed free-floating sections were mounted on Superfrost Plus slides (Menzel Gläser) and let dry overnight under fume hood. Hybridizations were performed as described by Abler et al.82 with modifications. After rehydration in PSBT, the sections were quenched at 25°C in 6% H_2_O_2_ (30 min) treated with proteinase K (10 min) and post-fixed with PFA-G (4% paraformaldehyde and 0.2% glutaraldehyde) for 20 min. Prehybridization step was performed in hybridization solution (HB: 50% formamide, 5XSSC, with 10μg/mL heparin, 10μg/mL yeast tRNA and, 1% Blocking reagent) for at least 1h at 60°C and then incubated overnight in HB with the digoxigenin-labeled riboprobes. Post-hybridization washes (50% formamide, 5x SSC, 1% SDS) were carried out for 2 hours at 60°C. Sections were washed in TNT (10mM Tris-HCL pH 7.5, 0.5M NaCl, 0.1% Tween™ 20) at 25°C and incubated for 15min at 37°C with RNase (0.25μg/mL), followed by a FS (50% formamide, 5x SSC) incubation of 2 hours (60°C). DIG was detected with an alkaline phosphatase labeled antibody (Roche).

After a saturation step in TBS pH 7.5 (10% sheep serum, 1% blocking reagent, 1% BSA, 0.1% Tween™ 20) for 1h (room temperature), sections were incubated overnight at 4°C with antibodies (1:1,000; in TBS containing: 5% sheep serum, 1% blocking reagent, 1% BSA). The following day, sections were washed for 2h in TBS (pH 7.5; 0.1% Tween™ 20 and 2mM levamisole) and then washed in alkaline phosphatase solution (100mM Tris-HCL pH 9.5, 100mM NaCl, 50mM MgCl_2_, 0.1% Tween™ 20 and 2mM levamisole). Bound antibodies were revealed using NBT-BCIP (Roche). After DIG in situ hybridization, slides were counterstained with DAPI (5 μg/mL, Invitrogen) washed and mounted using aqueous mounting.

### RTE-2_OV custom-made polyclonal antibodies

Custom-made polyclonal antibodies were obtained from Primm Biotech Custom Antibody Services (Milan, Italy) and raised against two peptides derived from two portions of the RTE-2_OV protein: GAA (1-100 aa), and HAA (569-673 aa) resulting to be unique within the translation of the assembled transcriptome. To choose the portion to select, the manufacturer also took into consideration: protein similarity (selection of regions with no significant identity to the murine and rabbit proteome); low complexity and transmembrane regions (exclusion of such regions); distribution of predicted antigenic peptides (selection of regions with a high number of predicted antigenic peptide). The selected synthetic peptides were injected into two rabbits and boosted three times within 38 days (at day: 21, 28, 35) after the first injection. The final bleeding was conducted 3 days after the last injection, and the crude sera were purified on Sepharose columns by immunoaffinity.

### Immunohistochemistry and antibody validation

SEM, SUB and OL dissected from *O. vulgaris* (N = 3) were immediately immersed 4% paraformaldehyde (PFA) in seawater (4°C for 3h). After fixation samples were washed several times in PBS, cryoprotected in sucrose 30% and embedded in OCT compound (OCT; Leica Biosystems). The embedded brain parts were then sectioned at 20 Am using a cryostat (Leica CM3050 S). No antigen retrieval was required. Tissue sections were rehydrated in three successive baths of 0.1 M PBS, and incubated for 90 min (at RT) in 5% goat serum (Vector Laboratories Ltd) diluted in 0.1 M PBS containing 0.05% Tween (PBTw). The slices were subsequently incubated at 4°C with custom-made polyclonal antibodies raised against LINE (G and H; see RTE-2_OV custom-made polyclonal antibodies for details). The next day, slices were again washed by several changes of PBTw and incubation (at RT for 90 min) with secondary antibodies was carried out using Alexa Fluor 488 or 546 goat anti-rabbit IgG both diluted 1:200 in PBTw. Subsequently, sections were rinsed and cell nuclei were counterstained with DAPI (Molecular Probes, Eugene, OR). Finally, after further extensive washes, sections were mounted with fluorescent mounting medium (Fluoromount, Sigma). For all antisera tested, omission of the primary antiserum and/or secondary antiserum resulted in negative staining. In addition, specificity was assured by pre-incubating (4°C, overnight) the antibodies with 1mg/mL of synthetic epitope (HAA and GAA, see RTE-2_OV custom-made polyclonal antibodies for details) before staining. Again, no immunostaining was observed. The two custom-made polyclonal antibodies raised against two different peptides of the RTE-2_OV protein stained the same spatial arrangement in the octopus brain tissue.

### Imaging

Sections were observed under microscopes depending on the techniqueImage acquisition and processing were performed using Leica Application Suite software (Leica Microsystems). For IHS we utilized a Leica DMI6000 B inverted microscope, and for IHC a Leica TCS SP8 X confocal microscope (Leica Microsystems, Germany). Tile Z-stacks were performed using a 0.2μm step size. IHC figures have been assembled following guidelines for color blindness provided by Wong [92].

## List of abbreviations

TEs: Transposable elements
LINE: Long interspersed element
SINE: Short interspersed element
LTR: Long terminal repeat
LncRNAs: Long non-coding RNAs
RTE: Retrotransposon element
SEM: Supra-esophageal mass
SUB: Sub-esophageal mass
OL: Optic lobe
ARM: Arm
GO: Gene ontology
CPM: Counts per million
RT-PCR: Reverse transcriptase-polymerase chain reaction
qPCR: Quantitative polymerase chain reaction
MELT: Mobile element locator tool
ISH: In-situ hybridization
IHC: Immunohistochemistry
RBP: Ribonucleoprotein

## Declarations

### Ethics approval and consent to participate

Samples were taken from local fishermen, by applying humane killing following principles detailed in Annex IV of Directive 2010/63/EU as described in the Guidelines on the Care and Welfare of Cephalopods (Fiorito et al. 2015) and following protocol for collection of tissues described by Baldascino and coworkers (Baldascino et al. 2017).

### Consent for publication

Not applicable.

### Availability of data and materials

*Octopus vulgaris* RNAseq raw reads have been submitted to the ArrayExpress database (www.ebi.ac.uk/arrayexpress) under accession number E-MTAB-3957. The data will be made public upon acceptance of this article. Login credential for referees are indicated in the first page of the supplementary pdf. All the remaining data are present in the supplementary files archive.

### Competing interests

The authors declare that they have no competing interests.

### Funding

The work has been supported by: Progetto Premiale MolEcOC (Italian Ministry of Education, University and Research, MIUR); Flagship project RITMARE (MIUR and Stazione Zoologica); BIOforIU PON Project (MIUR and European Regional Development Fund, FESR). Giuseppe Petrosino, Swaraj Basu, Massimiliano Volpe and Giulia Di Cristina have been supported by a SZN PhD fellowship.

## Authors Contributions

RS and GF conceived, designed and led the project; GPe, MV and RS performed bioinformatics analysis and interpreted the results with significant contributions from FM and SB; GPo carried out the in-situ hybridization and IHC experiments and choice sectioning and sampling of different animals; IZ cared of biological sampling and performed RNA extractions; SF and SG carried out the quantitative PCR experiments; DP and VB performed the RNAseq; OS and CA carried out extraction and sequencing of *O. bimaculoides* samples; IZ, CL, GDC, MTR, MIF and FR carried out the remaining molecular biology experiments and validations. RS and GF wrote the manuscript. All the Authors discussed the results and contributed to the manuscript.

## Acknowledgements

The authors would like to thank Paul Andrews for critical reading of the manuscript; Paolo Vatta and Marta Maurutto for assistance in CNV experiments; Marco Miralto and the BIOINforMA service at SZN for informatics support.

## Authors’ information: Current addresses

Giuseppe Petrosino: Institute of Molecular Biology (IMB), Mainz, Germany.

Francesco Musacchia: Central RNA Laboratory, Istituto Italiano di Tecnologia (IIT), Genova, Italy.

Swaray Basu: Strand Life Sciences, Bengaluru, India.

Giulia Di Cristina: Institute of Zoology, University of Cologne, Cologne, Germany.

Dinko Pavlinic: Institute of Molecular and Clinical Ophthalmology, Basel, Switzerland.

Massimiliano Volpe: Department of Biomedical and Clinical Sciences, Linköping University, Linköping, Sweden.

Oleg Simakov: Department of Molecular Evolution and Development, Wien University, Wien, Austria.

## Notes

### Competing Interest Statement

The authors have declared no competing interest.

